# Optimizing multiplexed imaging experimental design through tissue spatial segregation estimation

**DOI:** 10.1101/2021.11.28.470262

**Authors:** Pierre Bost, Daniel Schulz, Stefanie Engler, Clive Wasserfall, Bernd Bodenmiller

**Affiliations:** University of Zurich, Department of Quantitative Biomedicine, Zurich, 8057, Switzerland; ETH Zurich, Institute for Molecular Health Sciences, Zurich, 8093 Switzerland; Department of Pathology, Immunology, and Laboratory Medicine, Diabetes Institute, University of Florida, Gainesville, FL 32610, USA

## Abstract

Recent advances in multiplexed imaging methods allow simultaneous detection of dozens of proteins and hundreds of RNAs enabling deep spatial characterization of both healthy and diseased tissues. Parameters for design of optimal sequencing-based experiments have been established, but such parameters, especially those estimating how much area has to be imaged to capture all cell phenotype clusters, are lacking for multiplex imaging studies. Here, using a spatial transcriptomic atlas of healthy and tumor human tissues, we developed a new statistical framework that determines the number and area of fields of view necessary to accurately identify all cell types that are part of a tissue. Using this strategy on imaging mass cytometry data, we identified a measurement of tissue spatial segregation that enables optimal experimental design. This strategy will enable significantly improved design of multiplexed imaging studies.

## Main text

In the last decade, single-cell technologies for proteomic (Bendall et al. 2011), transcriptomic and genomic (Jaitin et al. 2014, Macosko et al. 2015, Xu et al 2012) analyses have been developed. Experiments using these technologies have enhanced our understanding of biological systems ranging from human immune cells (Villani et al. 2017) to whole cnidarian organisms (Sebé-Pedrós et al. 2018). Clear guidelines have been established to determine optimal experimental design of sequencing studies, including the total number of cells and sequencing depth necessary for detection of rare cell types or transcripts (Torre et al. 2018).

Increasingly, single-cell transcriptomic and proteomic measurements are performed with spatial resolution (Lewis et al. 2021). Multiplexed imaging techniques are modern counterparts of histological analyses and aim to detect a given set of cell types and their state based on the target markers used. Therefore, the ability of a multiplexed imaging experiment to detect every expected cell type that is present in a given tissue section is essential. Guidelines for optimal design of multiplexed imaging experiments, such as those performed using imaging mass cytometry (IMC) (Giesen et al. 2014), MIBI (Angelo et al. 2014), and CODEX (Goltsev et al. 2018) and in situ hybridization methods such as seqFISH (Shah et al. 2016), MERFISH (Chen et al. 2015) and HDST (Vickovic et al. 2019), have not been developed. Given that current highly-multiplexed tissue imaging methods have low spatial throughput and high costs, such guidelines, especially to estimate the area to be measured to capture a tissues phenotypic heterogeneity is urgently needed. Since it is possible to model the probability of detecting an object when imaging a given area (Illian et al. 2008), solid theoretical foundation for modeling and interpreting the outputs of multiplexed imaging experiments exists. Building on pioneering work on the number of regions that must be imaged to characterize the intensity distribution of a single fluorescent marker and single cell type (Rajaram et al. 2017), we report the development of a strategy to determine the minimal number of fields of view (FoVs) necessary to identify all main cell phenotypes across various healthy and tumor tissues.

### Optimal tissue sampling strategy for spatial transcriptomic data

Despite the lack of singe-cell resolution, spatial transcriptomic datasets cover large areas of tissues (16 mm^2^ for Visium® arrays) and are assumed to provide an exhaustive description of cell phenotypes present in the tissue. Using these spatial transcriptomic data, we wanted to assess how many areas have to be measured with another spatial imaging technology to capture all present phenotypes. We chose IMC which is capable to measure 40 markers in situ. We used previously collected 22 Visium datasets on 12 different types of tissues (Table S1). The Visium data were normalized and clustered to identify different cell types and cellular niches (Figure 1A). We then simulated IMC data acquisition on these same tissues by performing repeated random sampling without replacement of a variable number of non-overlapping small square regions with widths of 400 μm across the tissue. We computed the number of different clusters (which correspond to unique cell types) detected across the sampled regions. Finally, the results were aggregated across samplings. There was an apparent saturation in the detection of clusters as the number of FoVs increased (Figure 1B).

**Figure 1:**
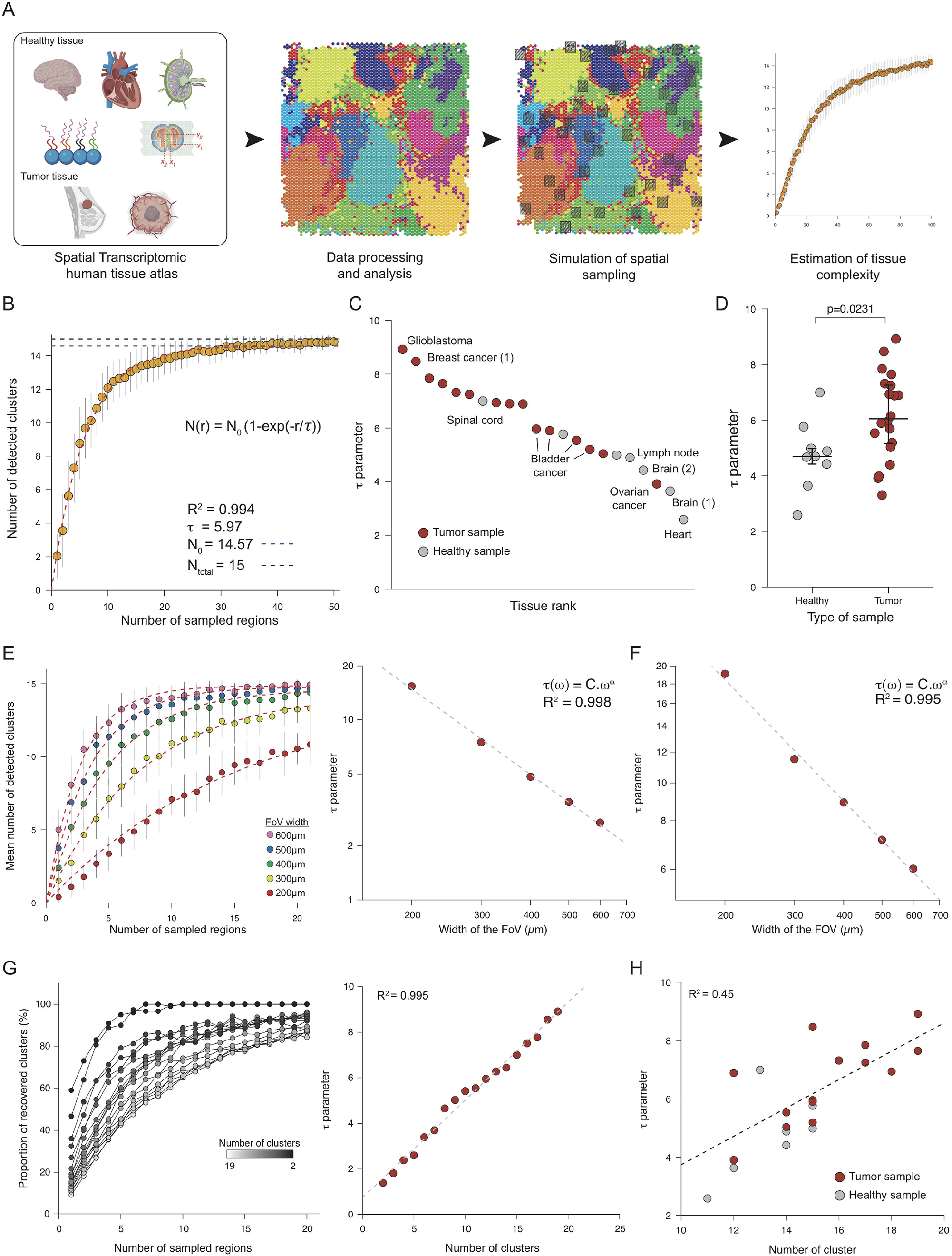
Use of spatial transcriptomic data to determine the optimal tissue sampling strategy for multiplexed imaging. **(A)** Analytical workflow used to simulate IMC of human tissues using spatial transcriptomic data. **(B)** Number of detected clusters vs. number of sampled regions for a bladder cancer Visium dataset with 400-μm FoVs. Each point corresponds to the mean number of recovered clusters across 50 similar simulations, and vertical bars correspond to standard error. The red dashed line corresponds to the fitted function. The horizontal dashed lines correspond to the total number of observed clusters (N_o_; blue) and the actual number of clusters (N_total_; grey). **(C)** Plot of *τ* for indicated samples from healthy and tumor samples. **(D)** Comparison of *τ* values from healthy and tumor samples. The p-value was computed using a Kruskal-Wallis rank test. **(E)** Left panel: Mean number of clusters detected vs. number of sampled regions for FoV widths ranging from 200 to 600 μm for the cerebellum Visium sample. Each point corresponds to the mean number of recovered cluster across 50 similar simulations, and vertical bars correspond to the standard error. Red dashed lines correspond to individual fits for each *w* value. Right panel: Relationship between *τ* and *w* for cerebellum sample. The dashed line corresponds to the linear regression after log10 transform. **(F)** Relationship between *τ* and *w* for the glioblastoma Visum sample. The dashed line corresponds to the linear regression after log10 transform. **(G)** Left panel: Proportion of clusters recovered as a function of *τ* for a glioblastoma sample for indicated number of clusters. Each point corresponds to the mean number of recovered clusters across 50 similar simulations. For the sake of clarity, the error bars and fitted curves are not displayed. Right panel: Relationship between *τ* and the number of clusters for a glioblastoma sample. The dashed line corresponds to a linear regression. **(H)** Relationship between *τ* and the number of clusters for all studied samples. The dashed line corresponds to a linear regression.

To model the relationship between number of clusters and number of FoVs, we used a model derived from the analysis of homogeneous Poisson point processes (Illian et al. 2008):

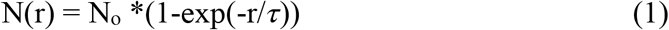

where *r* is the number of FoVs, N(r) is the number of recovered clusters, N_o_ corresponds to the total number of observed clusters, and *τ* indicates how many regions must be imaged to recover most of the known clusters. According to this model, 2*τ* FoVs must be imaged to detect 86% of known clusters. This model fit well across all Visium datasets (Figure S1A). The value of *τ* varied significantly across tissues (Figure 1C). We observed that tumor samples had higher *τ* values than healthy samples, indicating that more FoVs are required on average to identify cell phenotype clusters in tumor tissue than in healthy tissue (Figure 1D, p=0.0231).

We next studied the effects of the width of the FoV, *w*, on spatial sampling efficiency by performing the same simulated IMC analysis with various values of *w*. As expected, fewer regions needed to be imaged to detect all known clusters when *w* values were larger (Figure 1E, left panel). Following logarithmic transformation, there was a linear relationship between *w* and *τ* across all studied tissues (Figure 1E, right panel, Figure 1F, and Figure S1B). This indicated an underlying power law. Therefore, *τ* can be written as a function of *w*:

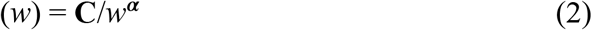

where **C** and ***α*** depend on sample.

We then explored whether there could be a relationship between *τ* and the granularity of the initial clustering analysis. To test this, we aggregated the most similar pairs of clusters for each sample by determining the correlation between mean expression profiles, performed a sampling analysis to compute *τ*, and then merged the next two most similar clusters, repeating until only two clusters were left (Figure S1C). We observed a linear relationship between *τ* and the number of clusters in certain Visium samples such as a glioblastoma (Figure 1G), but the linear regression fit poorly for others such as cerebellum (Figure S1D). In addition, across all samples, *τ* and the number of cell clusters were moderately correlated (Figure 1H, R^2^=0.45), implying that the number of clusters significantly impacts the value of *τ* and partly explains the difference in *τ* values between healthy and tumor samples. This indicates that this relationship cannot be generalized.

### Characterization of *α* and C as measures of tissue spatial segregation

In order to assess that results obtained from the analysis of spatial transcriptomic data can be generalized to data obtained from other technologies that provide images at single-cell resolution, we imaged large areas of two human formalin-fixed paraffin-embedded lymph node sections using IMC with two antibody panels (Table S2) and performed a spatial sampling analysis with various FoV widths (Figure 2A). As for the Visium lymph node data, the relationship between the number of sampled regions and of recovered clusters was fit by equation (1) (Figure 2B, left panel), and the FoV width affected *τ* as described by equation (2) (Figure 2B, middle and right panels). However, the values of *τ* drastically differed between the Visium lymph node data and the two IMC datasets (Figure 2C, left panel). When analyzing the parameters of (2), we found that the ***α*** parameter did not significantly differ between the two imaging modalities (Figure 2C, middle panel), whereas there was a large difference in **C** (Figure 2C, right panel). Within a given technology, the parameter ***α*** varied considerably across tissue types with values ranging from 2 for cardiac tissue to 0.91 for breast tumor tissue analyzed by spatial transcriptomics (Figure S2A). ***α*** was significantly lower in cancer samples than healthy tissues (Figure S2A, p=0.0401).

**Figure 2:**
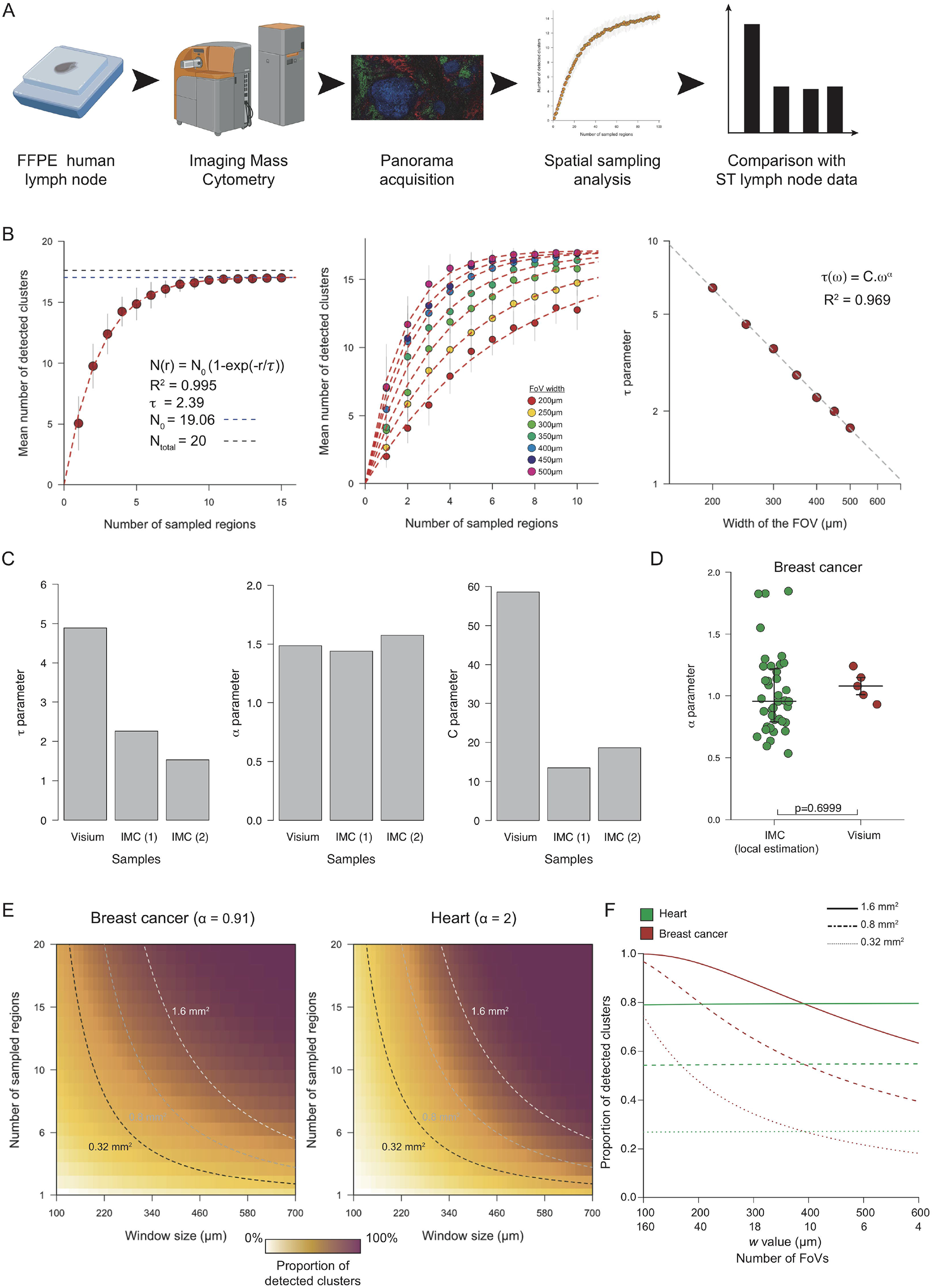
Identification of a technology invariant measure of tissue complexity. **(A)** Experimental workflow to compare the results of spatial transcriptomic and IMC large-scale analysis. **(B)** Left panel: Number of recovered clusters vs. number of sampled regions for IMC lymph node data. Each point corresponds to the mean number of recovered clusters across 50 similar simulations, and vertical bars correspond to the standard error. The red dashed line corresponds to the fitted function. The horizontal dashed lines correspond to number of observed clusters (N_o_; blue) and the actual number of clusters (N_total_; grey). Middle panel: Number of detected clusters vs. number of sampled regions for FoVs ranging from 200 to 500 μm for the IMC lymph node data from sample #1. Each point corresponds to the mean number of recovered cluster across 50 similar simulations, and vertical bars correspond to the standard error. The red dashed lines correspond to individual fits for each *w* value. Right panel: Relationship *τ* and *w* for the IMC lymph node data form sample #1. The dashed line corresponds to the linear regression after log10 transform. **(C)** Left panel: Values of *τ* for 400-μm width FoV for the Visium and IMC datasets for lymph node samples. Middle panel: Values of ***α*** for the lymph node datasets. Right panel: Values of **C** for the lymph node datasets. **(D)** Comparison of ***α*** values between the IMC breast cancer dataset and the five Visium breast cancer datasets. The p-value was computed using a Kruskall-Wallis test. **(E)** Estimation of sampling strategy efficiency for breast cancer (left panel) and heart (right panel) Visium samples. The dashed lines correspond to the possible values taken for a fixed area surface. **(F)** Proportions of detected clusters when area imaged (indicated by solid, dashed, or dotted lines) was fragmented for breast cancer (red lines) and heart (grey lines) datasets.

To further assess the properties of ***α***, we re-analyzed a previously published IMC dataset (Jackson et al. 2020) containing 100 FoVs, each derived from a unique breast cancer sample. For each FoV, we simulated a progressive shrinkage of the FoV width and computed the effect on the number of detected clusters to obtain an estimate of ***α*** for each FoV (Figure S2B). We did not observed a significant difference between the estimated ***α*** values using IMC data and the one estimated from five different Visium breast cancer samples (Figure 2D, p=0.699). These results support a hypothesis that ***α*** is a technology-independent but tissue dependent parameter.

Lastly, to evaluate the impact of ***α*** on the sampling strategy design, we computed the theoretical number of recovered clusters when sampling a defined area with various numbers and area of FoVs. We performed this analysis on two different types of tissue: cardiac (low spatial segregation, ***α***=2) and breast cancer tissue (strong spatial segregation, ***α***=0.91) using the values fitted on Visium datasets. First we observed that the number of recovered clusters was not affected by the fragmentation of the FoVs for the heart sample but only by the total imaged area (Figure 2E and 2F). In contrast, for breast cancer samples, increasing the fragmentation of the imaged area into multiple small ROIs drastically increases the number of recovered cell clusters: for instance when imaging 0.8mm^2^, shifting from 4 to 10 ROIs will result in the doubling of the proportion of recovered cell phenotypes, from less than 40% to more than 80% (Figure 2E and 2F).

## Discussion

Here we report how experimental design parameters impact the efficiency of multiplexed imaging experiments using the proportion of detected cell phenotypes as a simple yet robust metric. Our analysis identified the number of FoVs and their widths as key parameters that drive imaging experiment efficiency. Moreover, we determined the precise mathematical relationship linking these two parameters to the number of detected clusters. We found that the impact of FoV width on the experiment efficiency was regulated by a term ***α*** that seems to be tissue specific and potentially independent of the imaging technology used. In practice, ***α*** can be estimated in a pilot experiment using either a spatial transcriptomic approach, or by imaging a large region of a sample of a given cohort using a multiplexed imaging technology. Once determined for a tissue type, ***α*** should be valid for other tissues of similar type.

Interestingly, we observed highly variable values of ***α*** across tissues and this must be taken into account when planning an multiplexed imaging experiment. Indeed, a value of ***α*** close to 2 means that one can image a small number of large FoVs or many small FoVs and detect the same number of cell types. In contrast, a small ***α*** value requires the sampling of many small regions in order to efficiently recover the maximum number of cell phenotypes at a minimal cost. In order to facilitate the planning of imaging experiments, we provide ***α*** values for various healthy and cancerous tissues (Table S3).

Our model has limitations. In the moment, it only determines the ideal parameters to capture all (or a certain proportion) of phenotypic clusters present in the tissue. The model does not consider spatial relationships and tissues structures such as tertiary lymphoid structure. Additional work is therefore needed to see how our results, that are cell-based, can be extended to multi-cellular structures in order to implement more complex multiplex imaging experiments.

Beyond application to design of multiplexed imaging experiments, our results could also be used in the field of anapathology in which the current standard for classification of samples is the analysis of one to four circular punches of variable diameter (600 μm to 2 mm) (Eckel-Passow et al. 2010). Although we focused on the recovery of multiple cell types (i.e., clusters) rather than a single type of cell (for instance, HER2^+^ cells in breast cancer samples) (Harbeck et al. 2019), it is likely that a similar phenomenon of spatial segregation determines the efficacy of this type of sampling. In summary, our approach provides essential guidance for study of tissue structures using multiplex imaging in a time and cost-efficient manner.

## Methods

### Visium data pre-processing and analysis

Visium data were downloaded from the 10X Genomics website (support.10xgenomics.com/ spatial-gene-expression/datasets/), the Gene Expression Omnibus (GEO), or the Zenodo data portals. Spots with less than 1000 UMIs and genes with less than 100 UMIs were removed before any analysis. Data were analyzed by combining the classical single-cell RNA-seq pipeline **Pagoda2** (Lake et al. 2018) with a latent Dirichlet allocation analysis step. Briefly, the top 1500 most variable genes were identified using the adjustvariance() function from **Pagoda2** package, and the raw count data matrix containing only these genes was processed using the FitGoM() function from the **CountClust** package with a tolerance parameter set to 100 and the number of topics set to 5, 10, 15, or 20. For each number of topics, the BIC score was computed, and the number of topics displaying the lowest BIC or an elbow-like inflection was selected. The mixing matrix was then used for the next steps of analysis. A *k*-nearest-neighbor graph was built using the makeKnnGraph() function with parameter *k* set to 15 and using a cosine distance before performing a community detection analysis with the getKnnClusters() function with default parameters (corresponding to Louvain’s community detection (Blondel et al. 2008)).

### Spatial sampling analysis

To simulate spatial sampling strategies, we created a simple function that iteratively selects a random point on the sample, ‘draws’ a square with the sampled point at the center, and then checks whether this square overlaps with previously sampled squares. In case of overlap, the point is removed and a new point is sampled. A cluster was considered as detected by a given spatial sampling (set of sampled FoVs) if more than *T* spots belonging to that cluster were located in the drawn squares. The threshold *T* was changed based on the type of data: It was set to 2 for Visium data, 50 for the IMC lymph node data, and 20 for the IMC breast cancer data. This sampling was repeated 50 times to obtain a robust estimate.

The model proposed in equation (1) was derived from the analysis of a homogenous Poisson point (HPP) process defined by a density parameter ƛ. The probability that a random square of size *w* contain no points is equal to exp(-ƛ*w*^2^). A basic property of HPP processes is that the probability of finding no points in *r* independent (i.e., non-overlapping) squares is exp(-*ƛrw*^2^), and therefore the probability of finding at least one point is 1-exp(-*ƛnw*^2^). As we examined N_o_ different clusters (i.e., N_o_ different point processes), the mean number of detected clusters for a fixed number of squares N(r) is N_o_ =1- exp(-ƛ*nw*^2^), thus justifying the use of equation (1).

In order to fit the model described in equation (1), we used the nls() function with the following starting values: N_o_ of 20 and *τ* of 5. The quality of the fit was estimated using cor() and predict() functions. Fitting equation (2) to the data was done by first applying a log10 transform to the data before performing a classical least square regression using the lm() R function.

In order to study the effects of clustering granularity on cluster recovery, we first computed the mean expression of each gene in each cluster, then built a hierarchical clustering tree using Euclidean distance and Ward’s criterion. Then, using this tree, we iteratively merged the different clusters. At each step, we performed a spatial sampling analysis.

### Lymph node section processing and IMC data acquisition

The two lymph node formalin-fixed, paraffin-embedded blocks were first cut into 5-μm thick sections. They were then dewaxed and rehydrated and subjected to a heat-induced epitope retrieval step for 30 minutes at 95 ºC in 10 mM Tris, pH 9.2, 1 mM EDTA. The sections were then incubated in blocking buffer (3% BSA in TBS-T) for 1 hour at room temperature, before incubation with antibodies (diluted in blocking buffer) overnight at 4 ºC. Nuclear staining was then performed by adding an iridium solution (5 nM) diluted in TBS (1:100 dilution) to the sample and incubating for 5 minutes. The samples were then washed three times (10 minutes per wash) in TBS and dried. Images were acquired using an Hyperion Imaging System with the ablation frequency set to 200 Hz and the ablation energy set to 6 dB with X and Y steps set to 1 μm.

### IMC data pre-processing and analysis

The raw mcd files were processed using the Steinbock pipeline, version 0.70 (Windhager et al. 2021). In brief, the raw files were converted into tiff files, and the cells were segmented using a pre-trained neural network (Greenwald et al. 2021) using the H3K9ac channel as the nuclear channel and CD45RA/RO and Vimentin as the membrane channels. Default parameters were used for Mesmer with the exception of the —type parameter, which was set to ‘nuclei’. The mean channel intensity was then computed for each cell and exported as a text file, together with the location, the size, and other basic information on the cells. The single-cell IMC data were then analyzed using in-house R scripts (R version 4.0.3). Each channel was normalized by performing a Poisson regression between the total channel intensity and the cell size (in pixels); the Pearson’s residuals were extracted as the new scaled values. The cells were then clustered by first building a *k*-nearest-neighbor graph with 15 neighbors (using cosine distance) and then clustered using Louvain’s community detection implemented in the **igraph** package with default parameters.

### Breast cancer IMC data re-analysis

The SingleCellExperiment object containing single-cell information from 100 FoVs, each one derived from a different sample was downloaded from the Zenodo platform and analyzed using the following strategy: We first aggregated all cancer clusters (clusters 14 to 27) into a single cluster as the cancer clusters displayed a strong patient specificity. For each FoV, we progressively reduced the size of the image by factors of 1.1, 1.2, 1.5, 1.8, 2.2, 2.5, and 3 and computed the number of detected clusters. We then performed a linear regression between the log transformed number of detected clusters and the size of the reduced FoV using the lm() core function. FoVs with a low-quality model (R^2^ <0.9) were removed, and the slope of the regression was taken as the estimate of α. If we combine equations (1) and (2) when considering a single FoV, we have:

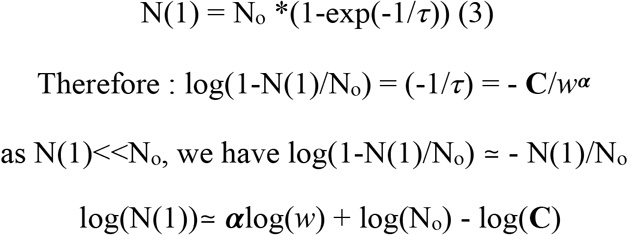

thus justifying our regression-based approach.

### Computing the effect of *α* on sampling strategy efficiency

In order to compute the number of recovered clusters in breast cancer and cardiac tissue as a function of both *r* (number of regions) and *w* (width of FoV), we substituted equation (2) into equation (1):

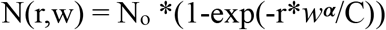

In order to compare both samples, we dropped the N_o_ term. We then selected three total area values (1.6 mm^2^, 0.8 mm^2^, and 0.32 mm^2^) and computed N(r,w) for different ratios of *r* and *w* with a constant r*w^2^ (total area) value.

## Author Contributions

P.B. developed the methodology, implemented the R code, performed the IMC experiments and wrote the manuscript. D.S. assisted with methodology development and implementation. S.E. performed IMC experiments. C.W. provided the samples and helped with methodology development. B.B. oversaw the project, assisted with study design, acquired funding, and helped write the manuscript.

## Acknowledgements

We acknowledge the Bodenmiller lab members for critical reading and providing feedback on the manuscript. P.B. is funded by an EMBO postdoctoral fellowship (fellowship number ALT 427-2021).

## Legends

**Supplementary Figure 1:**
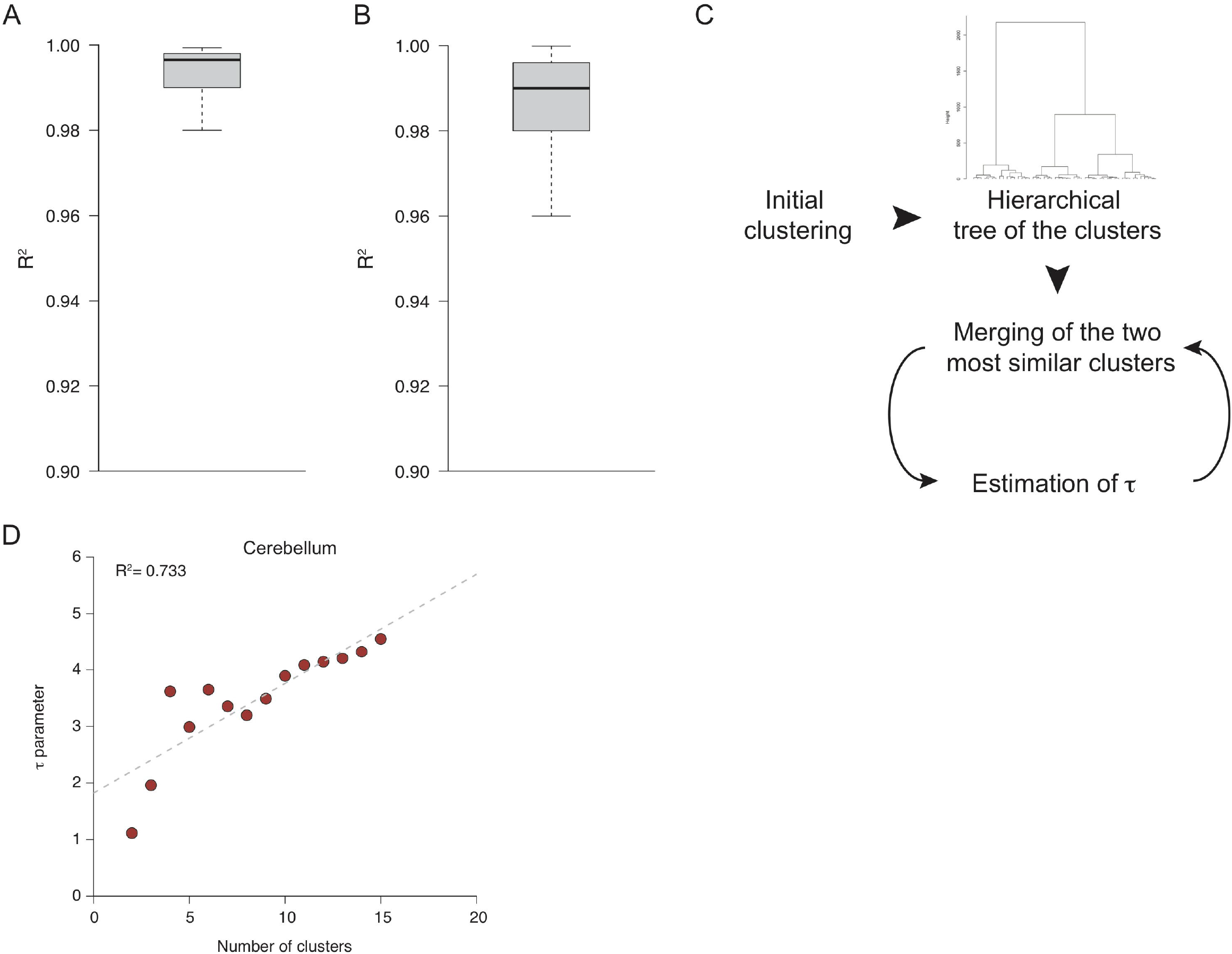
**(A)** Distribution of R^2^ values for the saturation model described in equation (1) across the Visium datasets. **(B)** Distribution of R^2^ values for the power-law model described in equation (2) across the Visium datasets. **(C)** Approach used to estimate the impact of clustering granularity on *τ*. **(D)** Relationship between *τ* and the number of clusters for the cerebellum sample.

**Supplementary Figure 2:**
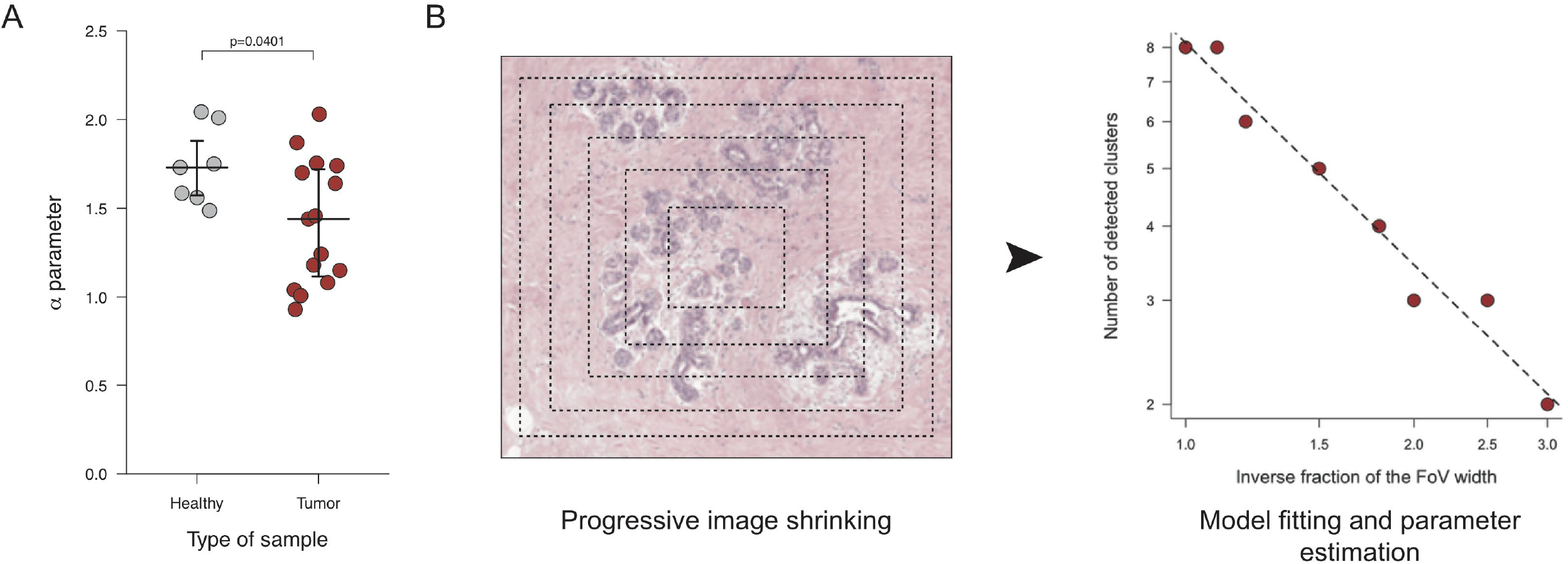
**(A)** Plot of ***α*** values from healthy and tumor samples. The p-value was computed using a Kruskal-Wallis rank test. **(B)** Approach used to estimate ***α*** from a set of small IMC FoVs.

